# Short sequence motif dynamics in the SARS-CoV-2 genome suggest a role for cytosine deamination in CpG reduction

**DOI:** 10.1101/2020.06.19.161687

**Authors:** Mukhtar Sadykov, Tobias Mourier, Qingtian Guan, Arnab Pain

## Abstract

RNA viruses use CpG reduction to evade the host cell defense, but the driving mechanisms are still largely unknown. In an attempt to address this we used a rapidly growing genomic dataset of SARS-CoV-2 with relevant metadata information. Remarkably, by simply ordering SARS-CoV-2 genomes by their date of collection, we find a progressive increase of C-to-U substitutions resulting in 5'-UCG-3' motif reduction that in turn have reduced the CpG frequency over just a few months of observation. This is consistent with APOBEC-mediated RNA editing resulting in CpG reduction, thus allowing the virus to escape ZAP-mediated RNA degradation. Our results thus link the dynamics of target sequences in the viral genome for two known host molecular defense mechanisms, mediated by the APOBEC and ZAP proteins.

---

Dear Editor,

The APOBEC protein family are host antiviral enzymes known for catalyzing cytosine to uracil deamination in foreign single-stranded DNA (ssDNA) and RNA (ssRNA) (Blanc and Davidson 2010; Salter and Smith 2018). Enzymatic target motifs for most of the APOBEC enzymes have been experimentally identified, among which the most common were 5′-[T/U]C-3′ and 5′-CC-3′ for DNA/RNA substrates (Salter and Smith 2018; McDaniel et al. 2020). It was recently suggested that the SARS-CoV-2 undergoes genome editing by host-dependent RNA-editing proteins such as APOBEC (Di Giorgio et al. 2020; Simmonds 2020; Rice et al. 2020; Schmidt et al. 2020).

Given the large amount of available data and the relatively low mutation rate of the SARS-CoV-2 virus (Rambaut et al. 2020), we aimed to monitor its genomic evolution on a very brief time scale during the COVID-19 pandemic. Here we demonstrate progressive C>U substitutions in SARS-CoV-2 genome within the timeframe of five months. We highlight the role of C>U substitutions in the reduction of 5′-UCG-3′ motifs and hypothesize that this progressive decrease is driven by host APOBEC activity.

We aligned 22,164 SARS-CoV-2 genomes from GISAID to the reference genome and observed a total of 9,210 single nucleotide changes with C>U being the most abundant (Figure 1A) (Figure S1 & S2; Table S1; Supplementary Text). Over a period of five months, we find a steady and substantial increase in C>U substitutions (Figure 1B), with almost half of them being synonymous (Supplementary Text, Figure S3), and not observed for other changes (Figure S4). One potential driver behind the increase in C>U changes could be the recently proposed APOBEC-mediated viral RNA editing (Di Giorgio et al. 2020; Simmonds 2020) (Supplementary Text). Since APOBEC3 family members display a preference for RNA in open conformation as opposed to forming secondary structures (McDaniel et al. 2020), we calculated the folding potential of all genomic sites that include C>U substitutions (Figure 1C). Positions with C>U changes are more often located in regions with low potential for forming secondary RNA structures. These observations are in agreement with the notion that members of the APOBEC family are the main drivers of cytosine deamination in SARS-CoV-2 (Di Giorgio et al. 2020; Simmonds 2020).

**Figure 1.**
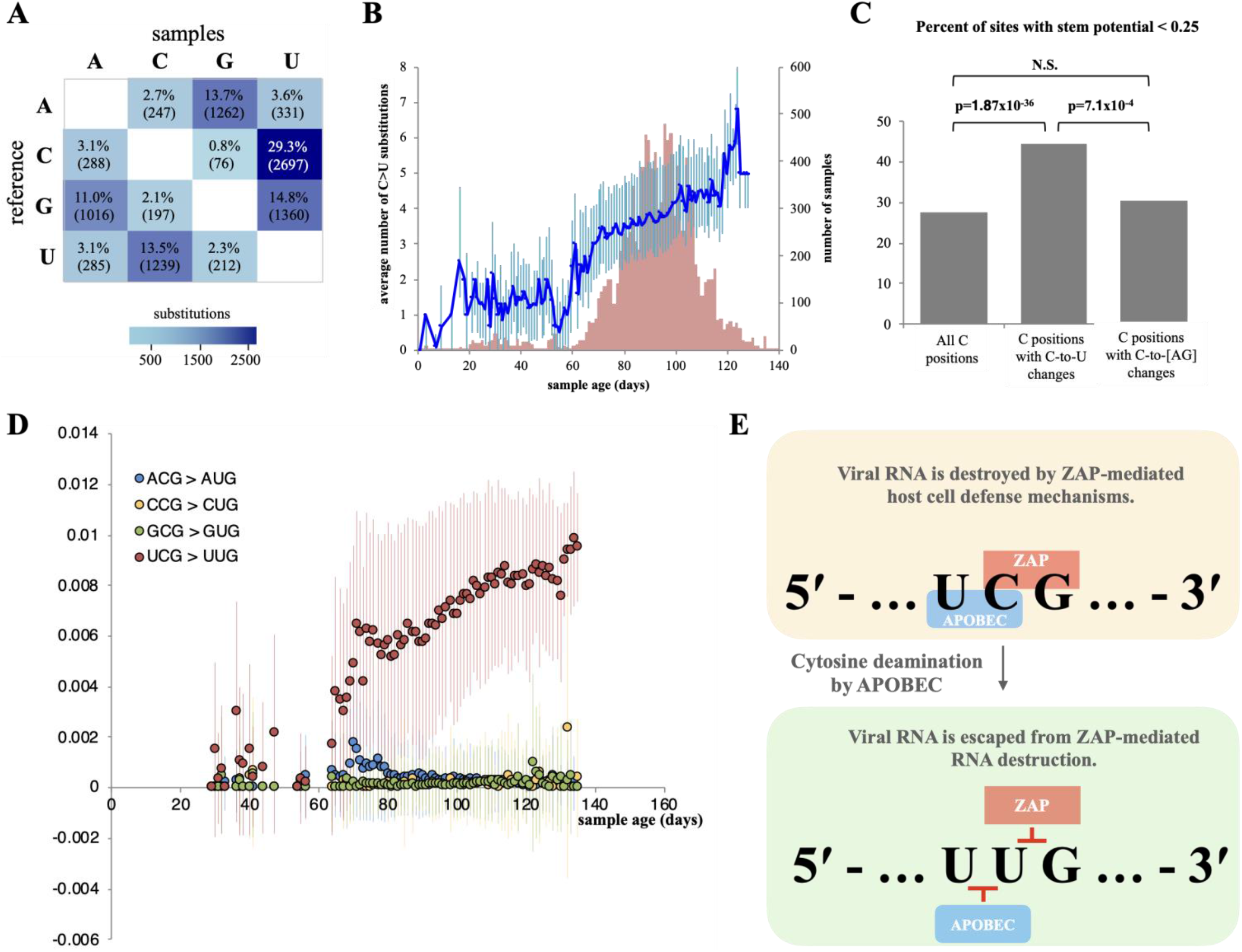
(A) SNV events observed between individual SARS-CoV-2 sample sequences (n=22,164) and the reference genome. (B) The number of C>U substitutions across sample dates. The average number of substitutions for each sampling day is plotted (blue line, left y-axis) with plus/minus one standard deviations as error bars. The number of samples for each day is shown as red bars (right y-axis). (C) Folding potential of positions with C>U changes (Supplementary Text). P-values from Fisher’s exact test are shown above bars. (D) The fraction of [A/C/G/U]CG triplets that are changed to [A/C/G/U]UG over time. The average fractions, relative to the reference genome, are shown as circles for each sampling day (x-axis). Error bars denote plus/minus one standard deviation. Only dates with at least 20 samples are plotted. (E) A model for the consequences of host-driven evolution by APOBEC enzymes on viral CpG dinucleotide composition.

We searched for possible APOBEC genetic footprints (5′-UC-3′ > 5′-UU-3′) in viral dinucleotide frequencies (Figure S5). Among all dinucleotides, UpC showed the highest degree of decrease, while UpU exerted the highest rates of increase, which is consistent with APOBEC activity (Supplementary Text).

When analyzing the context of genomic sites undergoing C>U changes, we noticed an enrichment for 5′-UCG-3′ motifs (Table S2). To assess the contribution of C>U changes in CpG loss, we examined the dynamics of [A/C/G/U]CG trinucleotides over time (Figure 1D). The progressive change (~1% over a 5-month period) of 5′-UCG-3′ to 5′-UUG-3′ is most striking when supported by a larger number of genomes (days 70 to 115), whereas no such pattern is observed for the other trinucleotides (Figure 1D). The association between cytosine deamination and CpG loss is further underlined by the rapid, progressive increase in 5′-UCG-3′ > 5′-UUG-3′ changes compared to other 5′-UC[A/C/U]-3′ motifs (Figure S7). No apparent progression of 5′-UCG-3′ over time is observed on the negative strand, suggesting that the action of APOBEC on the negative strand of SARS-CoV-2 is limited compared to the positive strand (Figure S8).

The zinc-finger antiviral protein (ZAP) selectively binds viral CpG regions that results in viral RNA degradation (Takata et al. 2017). Previous studies reported that the reduced number of CpG motifs in HIV and other viruses played an important role in the viral replication inside the host cell, allowing the virus to escape ZAP protein activity (Takata et al. 2017). Similarly, a strong suppression of CpGs is observed in SARS-CoV-2 compared to other coronaviruses (Digard et al. 2020). Given the high expression levels of APOBEC and ZAP genes in COVID-19 patients (Blanco-Melo et al. 2020), the direct interaction of APOBEC with viral RNA (Schmidt et al. 2020), and our observations, we hypothesize that as a consequence of APOBEC-mediated RNA editing, SARS-CoV-2 genome may escape host cell ZAP activity. Both APOBEC and ZAP are interferon-induced genes that act preferentially on ssRNA in open conformation (Luo et al. 2020; McDaniel et al. 2020). Initially, APOBEC and ZAP enzymes may have overlapping preferred target motifs for their enzymatic functions (Figure 1E). The catalytic activity of APOBEC on 5′-UC-3′ leads to cytosine deamination, which destroys ZAP’s specific acting site (5′-CG-3′). The conversion of C>U allows viral RNA to escape from ZAP-mediated RNA destruction. Therefore, uracil editing is more likely to become fixed at UCG positions due to the selective advantage this conveys to subvert ZAP-mediated degradation.

Our study of sequence dynamics across the SARS-CoV-2 pandemic supplements previous studies that by comparing the SARS-CoV-2 reference genome to other viral genomes address the evolutionary events prior to the Wuhan SARS-CoV-2 sequence. In contrast, our approach sheds light on the evolutionary events happening during the spread of SARS-CoV-2 among the human population.

A recent study hypothesized that both ZAP and APOBEC provide selective pressure that drives the adaptation of SARS-CoV-2 to its host (Wei et al. 2020). Here we provided one of the potential mechanisms that contribute to CpG reduction in SARS-CoV-2.

In summary, our phylogeny-free approach together with other recent studies strongly support the proposed model, and it merits future experimental validation. To our knowledge, this is the first study linking the dynamics of viral genome mutation to two known host molecular defense mechanisms, the APOBEC and ZAP proteins.

## Abbreviations

C>U: stands for cytosine to uracil substitution, the same applies to other nucleotide substitutions
APOBEC: Apolipoprotein B Editing Complex
ZAP: zinc-finger antiviral protein

## Acknowledgments

We thank all laboratories which have contributed sequences to the GISAID database, Zhadyra Yerkesh for giving her comments and helpful discussions.

This work was supported by funding from King Abdullah University of Science and Technology (KAUST) R3T initiative. Work in AP’s laboratory is supported by the KAUST faculty baseline fund (BAS/1/1020-01-01).

## Author Contributions

A.P. supervised the project. M.S. and T.M. designed experiments. T.M. and QG performed bioinformatic analysis. M.S. wrote the draft of the manuscript. All authors discussed, edited, read, and agreed to the final version of the manuscript.

## Availability of Data

The data underlying this article are available in GISAID, at https://gisaid.org. The ID numbers of genomes used are provided in Table S1.

## References

Blanc V, Davidson NO. 2010. APOBEC-1-mediated RNA editing. Wiley Interdiscip Rev Syst Biol Med. 2(5):594–602. doi:10.1002/wsbm.82.

Blanco-Melo D, Nilsson-Payant BE, Liu WC, Uhl S, Hoagland D, Møller R, Jordan TX, Oishi K, Panis M, Sachs D, et al. 2020. Imbalanced Host Response to SARS-CoV-2 Drives Development of COVID-19. Cell. 181(5):1036–1045.e9. doi:10.1016/j.cell.2020.04.026.

Digard P, Lee HM, Sharp C, Grey F, Gaunt E. 2020. Intra-genome variability in the dinucleotide composition of SARS-CoV-2. Virus Evol. 6(2). doi:10.1093/ve/veaa057.

Di Giorgio S, Martignano F, Torcia MG, Mattiuz G, Conticello SG. 2020. Evidence for host-dependent RNA editing in the transcriptome of SARS-CoV-2. Sci Adv. 6(25). doi:10.1126/sciadv.abb5813.

Luo X, Wang X, Gao Y, Zhu J, Liu S, Gao G, Gao P. 2020. Molecular Mechanism of RNA Recognition by Zinc-Finger Antiviral Protein. Cell Rep. 30(1):46–52.e4. doi:10.1016/j.celrep.2019.11.116.

McDaniel YZ, Wang D, Love RP, Adolph MB, Mohammadzadeh N, Chelico L, Mansky LM. 2020. Deamination hotspots among APOBEC3 family members are defined by both target site sequence context and ssDNA secondary structure. Nucleic Acids Res. 48(3):1353–1371. doi:10.1093/nar/gkz1164.

Rambaut A, Holmes EC, O’Toole Á, Hill V, McCrone JT, Ruis C, du Plessis L, Pybus OG. 2020. A dynamic nomenclature proposal for SARS-CoV-2 lineages to assist genomic epidemiology. Nat Microbiol. 5(11):1403–1407. doi:10.1038/s41564-020-0770-5.

Rice AM, Castillo Morales A, Ho AT, Mordstein C, Mühlhausen S, Watson S, Cano L, Young B, Kudla G, Hurst LD. 2020. Evidence for Strong Mutation Bias toward, and Selection against, U Content in SARS-CoV-2: Implications for Vaccine Design. Mol Biol Evol. doi:10.1093/molbev/msaa188.

Salter JD, Smith HC. 2018. Modeling the Embrace of a Mutator: APOBEC Selection of Nucleic Acid Ligands. Trends Biochem Sci. 43(8):606–622. doi:10.1016/j.tibs.2018.04.013.

Schmidt N, Lareau C, Keshishian H, Melanson R, Zimmer M, Kirschner L, Ade J, Werner S, Caliskan N, Lander E, et al. 2020. A direct RNA-protein interaction atlas of the SARS-CoV-2 RNA in infected human cells. bioRxiv.:2020.07.15.204404. doi:10.1101/2020.07.15.204404.

Simmonds P. 2020. Rampant C→U Hypermutation in the Genomes of SARS-CoV-2 and Other Coronaviruses: Causes and Consequences for Their Short- and Long-Term Evolutionary Trajectories. mSphere. 5(3). doi:10.1128/msphere.00408-20.

Takata MA, Gonçalves-Carneiro D, Zang TM, Soll SJ, York A, Blanco-Melo D, Bieniasz PD. 2017. CG dinucleotide suppression enables antiviral defence targeting non-self RNA. Nature. 550(7674):124–127. doi:10.1038/nature24039.

Wei Y, Silke J, Aris P, Xia X. 2020. Coronavirus genomes carry the signatures of their habitats. bioRxiv. doi:10.1101/2020.06.13.149591.

