## Supplementary Materials for "Short sequence motif dynamics in the SARS-CoV-2 genome suggest a role for cytosine deamination in CpG reduction"

**Supplementary Text**

**CpG dynamics in SARS-CoV-2**

Viruses utilize numerous mechanisms to avoid the host cell defense. One such mechanism is the CpG dinucleotide reduction observed in many single-stranded RNA viruses (Atkinson et al. 2014). Low CpG abundance is a consequence of viral mimicry of the host (Simmonds et al. 2013), that allows to escape selection pressure from diverse host cell defense mechanisms targeting CpG (Takata et al. 2017). Interestingly, a recent report showed the reduction of CpG dinucleotides in 5'-UTR region (Zhang et al. 2020). Our CpG net gain analysis showed a decline, indicating that the number of total CpG losses overrides their gain (Figure S5), in agreement with recently published observations (Wang et al. 2020; Zhang et al. 2020). Similar to other coronavirus genomes, SARS-CoV-2 has a low CpG frequency around 1.5%, while the expected frequency is 3.6% (Wang et al. 2020). The observed CpG decline could indicate the ongoing adaptation of the virus to the new host where antiviral proteins targeting foreign CpG regions are differently expressed (Xia 2020); and the new host acts as an evolutionary driver of CpG deselection (Digard et al. 2020). It was recently shown that synonymous substitutions disrupting CpG dinucleotides appear to contribute to the increased virulence of attenuated poliovirus (Stern et al. 2017). We observe that more than half of the C>U substitutions disrupting CpG dinucleotides were synonymous, consistent with recent studies (Di Gioacchino et al. 2020; Rice et al. 2020) (Figure S3). Similarly to our observation, the decline of CpGs upon zoonotic transfer into human host has been shown previously in the influenza A virus (Greenbaum et al. 2008). In the future, the SARS-CoV-2 virus may adapt its CpG frequency with its host, achieving so-called host-cell mimicry (Greenbaum et al. 2008), but the observed CpG loss suggests that equilibrium between gains and losses have not yet been reached.

**Summary of the approach**

Our time-series analyses are all carried out for each sample genome separately. Individual genomes are aligned to the reference genome and changes are recorded (see Methods). Only when plotting the progression over time, we are averaging over all samples from a given date.

Our analysis is centered around the temporal development after the Wuhan specimen was collected. All samples are defined by the amount of time between their collection date and the initial Wuhan reference.

In contrast to this, the number of changes presented in Figure 1A (main text) is found by counting the sites on the reference genome with a change in any sample. Hence, a change is counted only once regardless if it is found in all or just a single sample. This may lead to an underestimate of the most abundant changes (see next section).

**Probable underestimate of C>U frequency.**

As nearly half of the C positions in the reference are substituted to U in at least one sample, we expect a high degree of saturation. Downsizing the number of samples resulted in a consistent increase in C>U frequencies, suggesting that a considerable fraction of the observed C>U changes represent multiple, independent events (Figure S9). The reported C>U frequency is therefore most likely an underestimate.

**Comparison to previous studies**

Poulain and colleagues didn’t observe the evolutionary footprint of APOBEC-mediated editing in zoonotic coronaviruses using the K-mer representation ratio approach (Poulain et al. 2020). They discussed that the absence of an evolutionary footprint on the viral genome could be because of the relatively short duration of the pandemic and the presence of Vif-like protein encoded by ORF10. HIV genome contains Vif protein that upon binding to APOBEC leads to its further degradation, eventually avoiding the APOBEC restriction (Yu et al. 2003). In contrast to the K-mer representation counting approach, our analysis looks into the dynamics of short motifs according to the virus collection date. We show a gradual decrease of UpC and the increase of UpU dinucleotides over the first five months of viral evolution (Figure S5), indicating APOBEC footprinting in the SARS-CoV-2 genome.

Previously, from a comparative analysis of different coronaviruses, Woo and colleagues concluded that CpG reduction and cytosine deamination are two independent selective forces that shape coronavirus codon bias (Woo et al. 2007). Although selective forces may act differently upon motifs resulting from C>U changes, our results suggest a direct link between cytosine deamination and CpG reduction in SARS-CoV-2.

**Synonymous, non-synonymous and non-coding changes of C>U in CG and UCG sequence contexts.**

To better understand the C>U changes, we tested whether the substitutions were synonymous, non-synonymous or non-coding (Figure S3). The highest fraction of all C>U substitutions were synonymous. We observe that nearly 80% of the C>U substitutions disrupting CpG dinucleotides were synonymous, consistent with recent studies (Di Gioacchino et al. 2020; Rice et al. 2020). By further looking into C>U changes in UCG sequence context, similarly to CpG, the vast majority of changes were synonymous. The percentage of C>U changes in the non-coding regions were around 50-60%.

**Folding potential for UCG triplets**

Similarly to the analysis of folding potential for all C>U changes (see Methods and main Figure 1C), we tested for folding potential for UCG triplets with and without C>U changes in the middle position. 38 UCG triplets have UCG>UUG changes in at least five samples, 23 of these have stem-potential less than 0.25 (61%). Of 41 UCG triplets that have no UCG>UUG changes in any sample, 11 have stem-potential less than 0.25 (27%). Hence, UCG triplets with UCG>UUG changes display a significantly lower potential for RNA folding (Fisher's exact, p=0.0007832). However, when including UCG triplets that have UCG>UUG changes in less than 5 samples, this is no longer significant (p=0.5949).

**Regional differences in UCG changes**

High densities of UCG are located in the proximal and distal part of the viral genome, while the remaining genome (pos. 2,000-26,000 bp) show a roughly similar UCG density (Figure S6). Although СpG losses were previously noticed in the 5-UTR region (Zhang et al. 2020), our data suggests that UCG triplets are more often lost around the ORF1 region (pos. 266-21,555).

**Materials and Methods**

Samples and recording of substitutions

Samples were downloaded from GISAID (<https://www.gisaid.org/>) on May 23rd, 2020 (Elbe and Buckland-Merrett 2017). Only samples with available collection dates and with no more than 500 N's in the assembled sequence were used. December 24th, 2019 was set to day1. Assembled virus genomes were separately aligned to the reference genome (GenBank accession: NC_045512) using MAFFT (Katoh and Standley 2013), and mismatches were collected. Alignments with more than 500 gaps in the sample (likely due to incomplete assembly) and any alignments with gaps in the reference sequence were discarded. This resulted in 22,164 alignments. Only non-ambiguous bases and only mismatches without gaps in 3 positions up- and downstream were kept. To exclude alignment artifacts, mismatches in the first and last 100 positions in the reference genome were discarded.

The total number of substitutions is recorded from the combined set of pair-wise alignments, and a substitution at a given position in the reference is counted once regardless of the number of samples in which it is found. The timeline analysis of dinucleotide gains and losses were inferred from each sample's individual alignment to the reference.

Nucleotide motifs

For all di-, tri-, and tetra-nucleotide motifs containing C in the reference genome, the number of motifs with a C-to-U substitution, and the total number of the motifs were noted. The ratio between these two measures was compared to the expected ratio, defined as the number of C's with a C-to-U substitution divided by the total number of C's in the genome. The probability was calculated using a binomial distribution. All statistical tests were performed using Rstudio v. 1.1.414 (Lloyd and Sykes 1978).

Folding potential

The reference sequence was divided into overlapping 500-nucleotides windows, each shifted by 10 nucleotides. These sequences were folded using RNAfold (Lorenz et al. 2011). For each position in the reference genome, the number of windows in which the position was predicted to form a base-pair in a secondary structure was recorded. The folding potential of a position was defined as the fraction of windows with predicted base-pairing of that position. Hence, a position predicted to participate in a base-pair in all 500-nt windows has a folding potential of 1.

**Supplementary Figures and Tables**

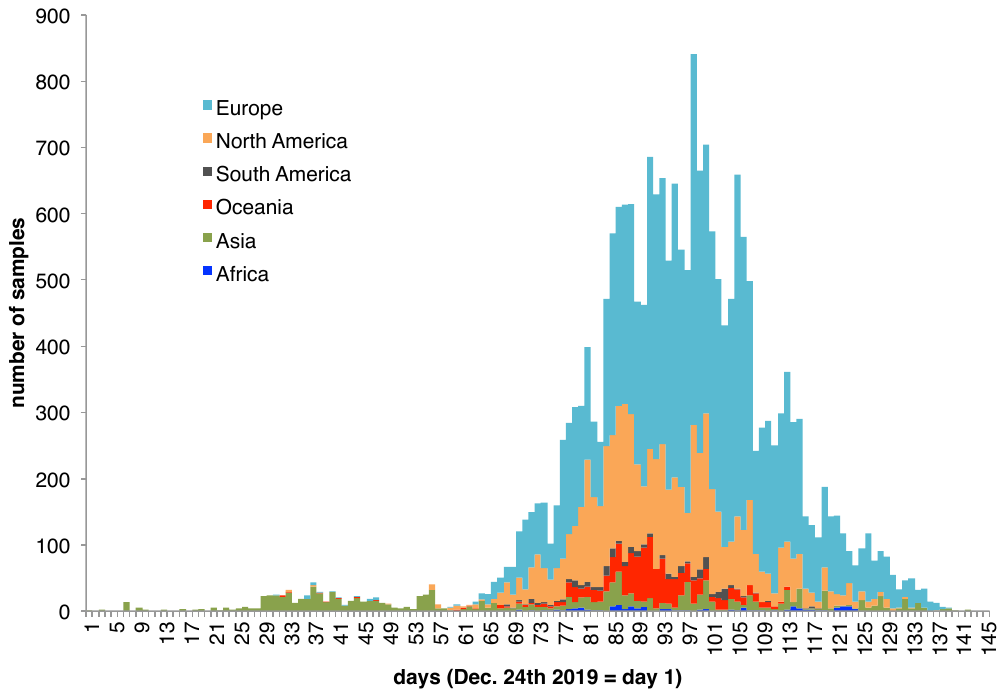

**Figure S1.** Number of samples from different time points. December 24^th^ 2019 is set to day 1. Samples are colored according to their geographic region.

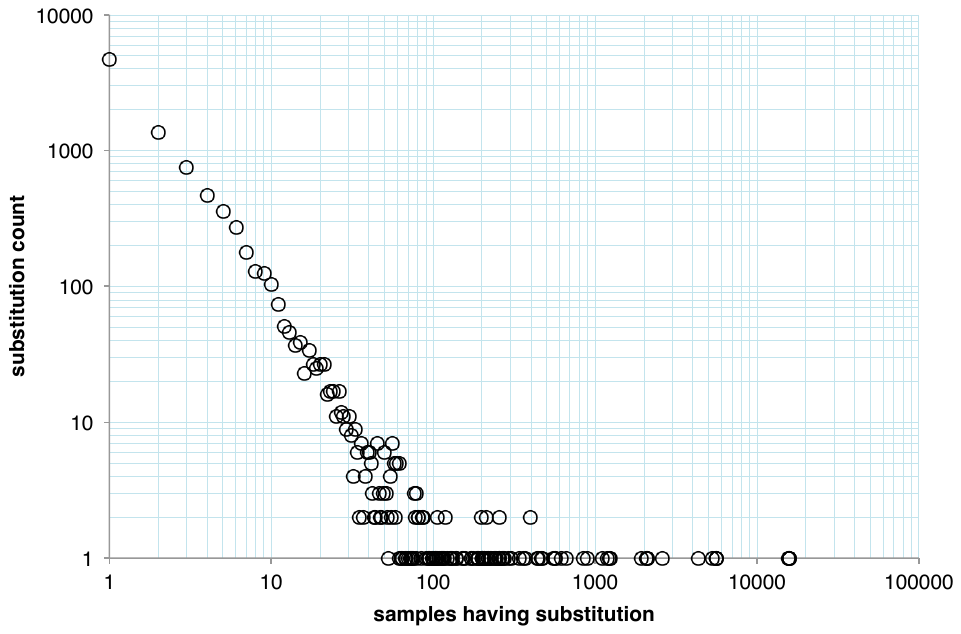

**Figure S2.** Substitution occurrences in samples. Most substitutions (4,683; top left) were observed in just a single sample (as denoted on the x-axis), whereas a single substitution was observed in 15,750 samples (bottom right).

**C>U**

**CG>UG**

**UCG>UUG**

Number of positions

Percentage of positions with changes

|  | non | C>T | %C>T |
| --- | --- | --- | --- |
| non-synonymous | 2174 | 1516 | 41.084 |
| synonymous | 556 | 1115 | 66.7265 |
| non-coding | 65 | 66 | 50.3817 |

Number of positions

Percentage of positions with changes

|  | non | C>T | %C>T |
| --- | --- | --- | --- |
| non-synonymous | 132 | 104 | 44.0678 |
| synonymous | 39 | 138 | 77.9661 |
| non-coding | 11 | 15 | 57.6923 |

Number of positions

Percentage of positions with changes

|  | non | C>T | %C>T |
| --- | --- | --- | --- |
| non-synonymous | 21 | 26 | 55.3191 |
| synonymous | 12 | 42 | 77.7778 |
| non-coding | 3 | 4 | 57.1429 |

**Figure S3.**

**Left:** Bars show the number of positions where a C-to-U change will be either non-synonymous, synonymous, or non-coding (as shown on the x-axis). Positions where a C-to-U change is observed are shown as blue parts of bars, and positions for which no change are shown in grey. The percentages of positions where C-to-U changes are observed are shown as open circles (right y-axis).

**Middle and Right:** Similar to the left plot, but only for C’s residing in CG dinucleotides and UCG trinucleotides, respectively.

average number of changes

age (days)

|  | A>C | A>G | A>U |
| --- | --- | --- | --- |
| 1 | 0 | 2 | 0 |
| 3 | 0 | 0 | 0 |
| 7 | 0.111111 | 0.444444 | 0 |
| 9 | 0.333333 | 0 | 0.333333 |
| 13 | 0 | 0 | 0 |
| 16 | 0 | 0 | 0 |
| 18 | 0 | 0 | 0 |
| 19 | 0 | 0 | 0 |
| 21 | 0.333333 | 1.33333 | 1 |
| 22 | 0 | 0 | 0 |
| 23 | 0 | 0 | 0 |
| 24 | 0 | 0 | 0 |
| 25 | 0 | 0 | 0 |
| 26 | 0 | 0 | 0 |
| 27 | 0 | 0 | 0 |
| 28 | 0 | 0.666667 | 0 |
| 29 | 0 | 0.1 | 0.1 |
| 30 | 0.0526316 | 0 | 0 |
| 31 | 0 | 0.411765 | 0.0588235 |
| 32 | 0.0454545 | 0.318182 | 0.0454545 |
| 33 | 0 | 0.0833333 | 0.0416667 |
| 34 | 0 | 0 | 0 |
| 35 | 0 | 0 | 0 |
| 36 | 0 | 0.217391 | 0.0434783 |
| 37 | 0.047619 | 0.285714 | 0.0238095 |
| 38 | 0.0357143 | 0.25 | 0 |
| 39 | 0 | 0.4 | 0.133333 |
| 40 | 0.0344828 | 0.206897 | 0.103448 |
| 41 | 0.05 | 0.25 | 0 |
| 42 | 0.111111 | 0.111111 | 0.111111 |
| 43 | 0.0666667 | 0.466667 | 0.2 |
| 44 | 0.0526316 | 0.315789 | 0 |
| 45 | 0 | 0.428571 | 0 |
| 46 | 0 | 0.333333 | 0.166667 |
| 47 | 0 | 0.166667 | 0 |
| 48 | 0 | 0.25 | 0 |
| 49 | 0 | 0.181818 | 0 |
| 50 | 0.2 | 0.4 | 0 |
| 51 | 0 | 0 | 0 |
| 52 | 0 | 0.333333 | 0 |
| 53 | 0 | 0 | 0 |
| 54 | 0 | 0.227273 | 0 |
| 55 | 0 | 0.12 | 0 |
| 56 | 0 | 0.025641 | 0.0512821 |
| 57 | 0 | 0.166667 | 0 |
| 58 | 0 | 0 | 0.25 |
| 59 | 0 | 0.6 | 0 |
| 60 | 0 | 0.375 | 0 |
| 61 | 0 | 0.428571 | 0.142857 |
| 62 | 0 | 0 | 0 |
| 63 | 0.0666667 | 0.733333 | 0.333333 |
| 64 | 0 | 0.576923 | 0.0384615 |
| 65 | 0 | 0.692308 | 0.0769231 |
| 66 | 0.0227273 | 0.545455 | 0 |
| 67 | 0 | 0.823529 | 0 |
| 68 | 0.0597015 | 0.820896 | 0 |
| 69 | 0.0769231 | 0.784615 | 0.0153846 |
| 70 | 0.0336134 | 0.857143 | 0.0504202 |
| 71 | 0.0147059 | 0.919118 | 0.0441176 |
| 72 | 0.0337838 | 0.918919 | 0.027027 |
| 73 | 0.0434783 | 0.962733 | 0.0559006 |
| 74 | 0.08125 | 0.96875 | 0.025 |
| 75 | 0.00980392 | 0.990196 | 0.0196078 |
| 76 | 0 | 0.981132 | 0.0188679 |
| 77 | 0.0540541 | 0.915058 | 0.046332 |
| 78 | 0.0248227 | 0.893617 | 0.0212766 |
| 79 | 0.0228013 | 0.973941 | 0.0944625 |
| 80 | 0.0392157 | 0.986928 | 0.0653595 |
| 81 | 0.0302267 | 1.02519 | 0.0277078 |
| 82 | 0.0282686 | 0.95053 | 0.0671378 |
| 83 | 0.0357143 | 1.12698 | 0.0357143 |
| 84 | 0.0191489 | 1.09362 | 0.0510638 |
| 85 | 0.024735 | 1.0265 | 0.0689046 |
| 86 | 0.0182119 | 1.06291 | 0.0546358 |
| 87 | 0.0212072 | 1.05057 | 0.040783 |
| 88 | 0.0392799 | 1.0311 | 0.0327332 |
| 89 | 0.0108932 | 0.980392 | 0.0348584 |
| 90 | 0.0196937 | 1 | 0.04814 |
| 91 | 0.017673 | 1 | 0.0265096 |
| 92 | 0.015949 | 1.03668 | 0.0350877 |
| 93 | 0.0168712 | 0.995399 | 0.0214724 |
| 94 | 0.0227273 | 1.02083 | 0.0189394 |
| 95 | 0.0201863 | 1.07453 | 0.0248447 |
| 96 | 0.0147874 | 1.07763 | 0.038817 |
| 97 | 0.0194932 | 1.03119 | 0.0331384 |
| 98 | 0.0228091 | 1.05762 | 0.0312125 |
| 99 | 0.0226244 | 1.05581 | 0.0769231 |
| 100 | 0.031339 | 1.03704 | 0.034188 |
| 101 | 0.0174825 | 1.06469 | 0.0367133 |
| 102 | 0.026 | 1.084 | 0.022 |
| 103 | 0.0255814 | 1.08837 | 0.0372093 |
| 104 | 0.0127119 | 1.10381 | 0.0402542 |
| 105 | 0.0182927 | 1.14939 | 0.035061 |
| 106 | 0.0283186 | 1.11858 | 0.0283186 |
| 107 | 0.0221328 | 1.10664 | 0.0221328 |
| 108 | 0.0371901 | 1.06198 | 0.0206612 |
| 109 | 0.0252708 | 1.16245 | 0.00722022 |
| 110 | 0.0313589 | 1.07317 | 0.0383275 |
| 111 | 0.0281124 | 1.22088 | 0.0281124 |
| 112 | 0.0402685 | 1.22819 | 0.0436242 |
| 113 | 0.0360111 | 1.19668 | 0.0360111 |
| 114 | 0.0385965 | 1.24912 | 0.0245614 |
| 115 | 0.0206897 | 1.1931 | 0.0310345 |
| 116 | 0.034965 | 1.23077 | 0.041958 |
| 117 | 0.0230769 | 1.16923 | 0.0307692 |
| 118 | 0.00900901 | 1.30631 | 0.027027 |
| 119 | 0.0106952 | 1.15508 | 0.0160428 |
| 120 | 0.0979021 | 1.21678 | 0.0629371 |
| 121 | 0.0555556 | 1.40278 | 0.0347222 |
| 122 | 0.0169492 | 1.31356 | 0.0762712 |
| 123 | 0.032967 | 1.18681 | 0.043956 |
| 124 | 0.057971 | 1.30435 | 0.0724638 |
| 125 | 0.0315789 | 1.31579 | 0.0421053 |
| 126 | 0.042735 | 1.30769 | 0.017094 |
| 127 | 0.0526316 | 1.34211 | 0.0921053 |
| 128 | 0.0769231 | 1.34066 | 0.10989 |
| 129 | 0.0853659 | 1.43902 | 0.0731707 |
| 130 | 0.0363636 | 1.30909 | 0.0727273 |
| 131 | 0.0606061 | 1.48485 | 0.030303 |
| 132 | 0.0217391 | 1.26087 | 0.0217391 |
| 133 | 0.02 | 1.66 | 0.08 |
| 134 | 0.125 | 1.21875 | 0.09375 |
| 135 | 0 | 1.55882 | 0.117647 |
| 136 | 0 | 1.4 | 0 |
| 137 | 0.0769231 | 1.61538 | 0.153846 |
| 138 | 0.166667 | 1.33333 | 0 |
| 139 | 0 | 1.4 | 0 |
| 140 | 0 | 0 | 0 |
| 142 | 0 | 1.5 | 0.5 |

|  | C>A | C>G | C>U |
| --- | --- | --- | --- |
| 1 | 0 | 0 | 0 |
| 3 | 1 | 0 | 0 |
| 7 | 0 | 0.111111 | 0.111111 |
| 9 | 0 | 0 | 0.666667 |
| 13 | 0 | 0 | 1 |
| 16 | 0 | 0 | 4 |
| 18 | 0 | 0 | 1.5 |
| 19 | 0 | 0 | 1 |
| 21 | 0 | 0.333333 | 2 |
| 22 | 0 | 0 | 2 |
| 23 | 0 | 0 | 1.5 |
| 24 | 0 | 0 | 1 |
| 25 | 0.333333 | 0 | 1.33333 |
| 26 | 0 | 0 | 1.16667 |
| 27 | 0 | 0.5 | 2 |
| 28 | 0 | 0 | 0.666667 |
| 29 | 0 | 0 | 1.9 |
| 30 | 0.0526316 | 0 | 1.57895 |
| 31 | 0.0588235 | 0 | 1.05882 |
| 32 | 0 | 0.0454545 | 1.36364 |
| 33 | 0.0833333 | 0 | 1.04167 |
| 34 | 0 | 0 | 1.23077 |
| 35 | 0.0588235 | 0.0588235 | 1.05882 |
| 36 | 0.0434783 | 0.0434783 | 1.6087 |
| 37 | 0.0238095 | 0.0238095 | 1.64286 |
| 38 | 0 | 0 | 1.42857 |
| 39 | 0.0666667 | 0 | 1.66667 |
| 40 | 0 | 0.0344828 | 1.51724 |
| 41 | 0.05 | 0 | 1.55 |
| 42 | 0 | 0 | 1.55556 |
| 43 | 0.0666667 | 0 | 1.53333 |
| 44 | 0 | 0 | 1.21053 |
| 45 | 0.0714286 | 0 | 1.21429 |
| 46 | 0 | 0.0555556 | 1.44444 |
| 47 | 0 | 0.166667 | 1.94444 |
| 48 | 0.0833333 | 0 | 1.25 |
| 49 | 0 | 0 | 1.54545 |
| 50 | 0 | 0 | 2 |
| 51 | 0 | 0 | 1.75 |
| 52 | 0 | 0 | 1.16667 |
| 53 | 0 | 0 | 0.666667 |
| 54 | 0 | 0 | 0.727273 |
| 55 | 0 | 0 | 0.52 |
| 56 | 0.0769231 | 0 | 0.820513 |
| 57 | 0 | 0 | 1.16667 |
| 58 | 0 | 0 | 1 |
| 59 | 0 | 0 | 1.6 |
| 60 | 0 | 0 | 2.375 |
| 61 | 0 | 0.142857 | 2.42857 |
| 62 | 0 | 0 | 1.57143 |
| 63 | 0.333333 | 0 | 2.33333 |
| 64 | 0.0384615 | 0.0384615 | 2.30769 |
| 65 | 0 | 0 | 2.42308 |
| 66 | 0.0454545 | 0 | 2.20455 |
| 67 | 0.0588235 | 0 | 2.72549 |
| 68 | 0 | 0 | 3.02985 |
| 69 | 0.0153846 | 0 | 2.95385 |
| 70 | 0.0672269 | 0.00840336 | 3.17647 |
| 71 | 0.0514706 | 0.00735294 | 3.38971 |
| 72 | 0.0202703 | 0.00675676 | 3.43919 |
| 73 | 0.0496894 | 0 | 3.21739 |
| 74 | 0.05 | 0.00625 | 3.25 |
| 75 | 0.0588235 | 0.00980392 | 3.26471 |
| 76 | 0.0377358 | 0 | 3.42767 |
| 77 | 0.0501931 | 0.00772201 | 3.52896 |
| 78 | 0.0496454 | 0.0035461 | 3.35106 |
| 79 | 0.0781759 | 0.0130293 | 3.30945 |
| 80 | 0.0588235 | 0 | 3.53595 |
| 81 | 0.0528967 | 0.00251889 | 3.40554 |
| 82 | 0.0212014 | 0.00353357 | 3.46643 |
| 83 | 0.0119048 | 0.00793651 | 3.59921 |
| 84 | 0.0361702 | 0.0106383 | 3.54255 |
| 85 | 0.0441696 | 0.0141343 | 3.65018 |
| 86 | 0.0546358 | 0.00827815 | 3.83444 |
| 87 | 0.0489396 | 0.0163132 | 3.71289 |
| 88 | 0.091653 | 0.0196399 | 3.58265 |
| 89 | 0.0413943 | 0.0196078 | 3.71895 |
| 90 | 0.0568928 | 0.00875274 | 3.64114 |
| 91 | 0.0382916 | 0.0103093 | 3.67894 |
| 92 | 0.0526316 | 0.0143541 | 3.66667 |
| 93 | 0.0582822 | 0.00766871 | 3.67945 |
| 94 | 0.0662879 | 0.00568182 | 3.81439 |
| 95 | 0.068323 | 0.0139752 | 3.81677 |
| 96 | 0.0702403 | 0.0166359 | 3.86506 |
| 97 | 0.0779727 | 0.00779727 | 3.88499 |
| 98 | 0.0456182 | 0.0060024 | 4.01681 |
| 99 | 0.0965309 | 0.0120664 | 4.04827 |
| 100 | 0.102564 | 0.00569801 | 3.95299 |
| 101 | 0.0821678 | 0.0122378 | 4.1049 |
| 102 | 0.048 | 0.004 | 4.172 |
| 103 | 0.0697674 | 0.00465116 | 4.0186 |
| 104 | 0.0699153 | 0.00211864 | 3.92161 |
| 105 | 0.0564024 | 0.00762195 | 4.16463 |
| 106 | 0.0389381 | 0.00530973 | 4.07611 |
| 107 | 0.128773 | 0.00402414 | 4.22736 |
| 108 | 0.0867769 | 0.0123967 | 3.8843 |
| 109 | 0.0397112 | 0.0108303 | 4.17329 |
| 110 | 0.0731707 | 0.00348432 | 4.38676 |
| 111 | 0.0682731 | 0.0240964 | 4.15261 |
| 112 | 0.100671 | 0.0100671 | 4.45638 |
| 113 | 0.099723 | 0.0138504 | 4.42105 |
| 114 | 0.0912281 | 0.0105263 | 4.58947 |
| 115 | 0.0793103 | 0.00689655 | 4.22759 |
| 116 | 0.0629371 | 0.013986 | 4.41259 |
| 117 | 0.0846154 | 0 | 4.7 |
| 118 | 0.0630631 | 0 | 4.26126 |
| 119 | 0.0855615 | 0 | 4.57219 |
| 120 | 0.202797 | 0.00699301 | 4.18881 |
| 121 | 0.131944 | 0 | 4.26389 |
| 122 | 0.110169 | 0 | 4.62712 |
| 123 | 0.197802 | 0 | 5 |
| 124 | 0.217391 | 0 | 4.6087 |
| 125 | 0.178947 | 0.0105263 | 4.93684 |
| 126 | 0.17094 | 0 | 4.59829 |
| 127 | 0.0921053 | 0.0131579 | 5.05263 |
| 128 | 0.032967 | 0.010989 | 4.84615 |
| 129 | 0.0243902 | 0.0243902 | 4.63415 |
| 130 | 0.0363636 | 0 | 4.16364 |
| 131 | 0.030303 | 0 | 4.9697 |
| 132 | 0 | 0 | 5.34783 |
| 133 | 0.02 | 0 | 4.46 |
| 134 | 0.0625 | 0 | 5.09375 |
| 135 | 0.0294118 | 0 | 4.67647 |
| 136 | 0.133333 | 0 | 5.33333 |
| 137 | 0 | 0 | 4.69231 |
| 138 | 0.166667 | 0 | 5.66667 |
| 139 | 0.4 | 0 | 5.4 |
| 140 | 2 | 0 | 6 |
| 142 | 0 | 0 | 3.5 |

|  | G>A | G>C | G>U |
| --- | --- | --- | --- |
| 1 | 0 | 0 | 0 |
| 3 | 0 | 0 | 0 |
| 7 | 0.555556 | 0 | 0 |
| 9 | 0 | 0 | 0.333333 |
| 13 | 0 | 0 | 0 |
| 16 | 1 | 0 | 0 |
| 18 | 0.5 | 0 | 0 |
| 19 | 0.666667 | 0 | 0 |
| 21 | 0 | 1 | 0.333333 |
| 22 | 0 | 0 | 0 |
| 23 | 0 | 0.25 | 0 |
| 24 | 0 | 0 | 0 |
| 25 | 0 | 0 | 0.333333 |
| 26 | 0.166667 | 0.333333 | 0.666667 |
| 27 | 0.5 | 0 | 1 |
| 28 | 0 | 0 | 0.666667 |
| 29 | 0 | 0.2 | 0.4 |
| 30 | 0.105263 | 0.157895 | 0.473684 |
| 31 | 0.117647 | 0.176471 | 0.823529 |
| 32 | 0.272727 | 0.272727 | 0.5 |
| 33 | 0.166667 | 0.0416667 | 0.375 |
| 34 | 0.230769 | 0 | 0.769231 |
| 35 | 0.0588235 | 0 | 0.470588 |
| 36 | 0.173913 | 0.0434783 | 0.521739 |
| 37 | 0.166667 | 0 | 0.642857 |
| 38 | 0.428571 | 0 | 0.535714 |
| 39 | 0.2 | 0 | 0.466667 |
| 40 | 0.103448 | 0.137931 | 0.172414 |
| 41 | 0.3 | 0.05 | 0.4 |
| 42 | 0.333333 | 0 | 0.555556 |
| 43 | 0.133333 | 0 | 0.533333 |
| 44 | 0.368421 | 0 | 0.263158 |
| 45 | 0.357143 | 0 | 0.857143 |
| 46 | 0.277778 | 0 | 0.611111 |
| 47 | 0.277778 | 0.111111 | 0.277778 |
| 48 | 0.0833333 | 0.166667 | 0.916667 |
| 49 | 0 | 0 | 1.09091 |
| 50 | 0.2 | 0.2 | 0.8 |
| 51 | 0.5 | 0.25 | 0.5 |
| 52 | 0.166667 | 0 | 0.833333 |
| 53 | 0 | 0 | 1 |
| 54 | 0.318182 | 0.0454545 | 1.27273 |
| 55 | 0.16 | 0.04 | 1.16 |
| 56 | 0.102564 | 0 | 1.05128 |
| 57 | 0 | 0.166667 | 2.33333 |
| 58 | 0.25 | 0 | 2.25 |
| 59 | 0.2 | 0 | 0.6 |
| 60 | 0.125 | 0.125 | 0.75 |
| 61 | 0.142857 | 0 | 1 |
| 62 | 0.285714 | 0.142857 | 0.714286 |
| 63 | 0.4 | 0.133333 | 0.533333 |
| 64 | 0.692308 | 0.115385 | 0.923077 |
| 65 | 0.692308 | 0.115385 | 0.884615 |
| 66 | 0.909091 | 0.227273 | 0.909091 |
| 67 | 0.666667 | 0.176471 | 0.568627 |
| 68 | 0.447761 | 0.164179 | 0.656716 |
| 69 | 0.430769 | 0.215385 | 0.630769 |
| 70 | 0.781513 | 0.336134 | 0.495798 |
| 71 | 0.889706 | 0.375 | 0.375 |
| 72 | 0.641892 | 0.243243 | 0.621622 |
| 73 | 0.453416 | 0.26087 | 0.459627 |
| 74 | 0.59375 | 0.30625 | 0.55625 |
| 75 | 0.539216 | 0.245098 | 0.666667 |
| 76 | 0.515723 | 0.207547 | 0.63522 |
| 77 | 0.490347 | 0.200772 | 0.733591 |
| 78 | 0.542553 | 0.205674 | 0.797872 |
| 79 | 0.592834 | 0.224756 | 0.62215 |
| 80 | 0.486928 | 0.189542 | 0.712418 |
| 81 | 0.448363 | 0.15869 | 0.717884 |
| 82 | 0.568905 | 0.173145 | 0.862191 |
| 83 | 0.424603 | 0.130952 | 0.912698 |
| 84 | 0.491489 | 0.180851 | 0.795745 |
| 85 | 0.446996 | 0.19258 | 0.90636 |
| 86 | 0.524834 | 0.200331 | 0.915563 |
| 87 | 0.49429 | 0.195759 | 0.91354 |
| 88 | 0.600655 | 0.227496 | 0.854337 |
| 89 | 0.655773 | 0.250545 | 0.825708 |
| 90 | 0.689278 | 0.245077 | 0.881838 |
| 91 | 0.661267 | 0.228277 | 1.01473 |
| 92 | 0.679426 | 0.261563 | 0.987241 |
| 93 | 0.616564 | 0.239264 | 0.894172 |
| 94 | 0.657197 | 0.253788 | 0.837121 |
| 95 | 0.726708 | 0.287267 | 0.82764 |
| 96 | 0.691312 | 0.277264 | 0.787431 |
| 97 | 0.766082 | 0.309942 | 0.808967 |
| 98 | 0.776711 | 0.284514 | 0.889556 |
| 99 | 0.773756 | 0.307692 | 0.825038 |
| 100 | 0.739316 | 0.270655 | 0.867521 |
| 101 | 0.828671 | 0.332168 | 0.933566 |
| 102 | 0.832 | 0.322 | 0.908 |
| 103 | 1.04419 | 0.404651 | 0.795349 |
| 104 | 1.03814 | 0.455508 | 0.815678 |
| 105 | 1.00915 | 0.419207 | 0.815549 |
| 106 | 1.13274 | 0.460177 | 0.787611 |
| 107 | 1 | 0.402414 | 0.847082 |
| 108 | 0.92562 | 0.371901 | 0.698347 |
| 109 | 1.05776 | 0.418773 | 0.960289 |
| 110 | 1.27178 | 0.519164 | 0.721254 |
| 111 | 1.15261 | 0.493976 | 0.851406 |
| 112 | 0.865772 | 0.348993 | 0.909396 |
| 113 | 0.969529 | 0.421053 | 0.911357 |
| 114 | 1.09474 | 0.442105 | 0.712281 |
| 115 | 1.00345 | 0.413793 | 0.789655 |
| 116 | 1.1958 | 0.440559 | 0.839161 |
| 117 | 1.03846 | 0.4 | 0.761538 |
| 118 | 1.18919 | 0.513514 | 0.738739 |
| 119 | 0.860963 | 0.342246 | 0.957219 |
| 120 | 0.923077 | 0.391608 | 0.671329 |
| 121 | 0.9375 | 0.361111 | 0.8125 |
| 122 | 0.855932 | 0.398305 | 0.694915 |
| 123 | 0.483516 | 0.208791 | 1.06593 |
| 124 | 1.01449 | 0.318841 | 0.73913 |
| 125 | 0.747368 | 0.315789 | 0.989474 |
| 126 | 1.05983 | 0.521368 | 0.726496 |
| 127 | 1.23684 | 0.5 | 0.921053 |
| 128 | 1.10989 | 0.43956 | 0.802198 |
| 129 | 1.15854 | 0.487805 | 0.829268 |
| 130 | 1.41818 | 0.581818 | 0.545455 |
| 131 | 1.06061 | 0.424242 | 1.0303 |
| 132 | 1.04348 | 0.304348 | 1.15217 |
| 133 | 0.86 | 0.36 | 0.84 |
| 134 | 1.03125 | 0.40625 | 1.3125 |
| 135 | 1.41176 | 0.588235 | 0.882353 |
| 136 | 1.33333 | 0.6 | 0.866667 |
| 137 | 1.23077 | 0.461538 | 1.30769 |
| 138 | 0.333333 | 0.166667 | 0.833333 |
| 139 | 1.4 | 0.4 | 1.6 |
| 140 | 1 | 0 | 3 |
| 142 | 2 | 0.5 | 2 |

|  | U>A | U>C | U>G |
| --- | --- | --- | --- |
| 1 | 1 | 0 | 0 |
| 3 | 1 | 0 | 0 |
| 7 | 0.222222 | 0.333333 | 0.111111 |
| 9 | 0 | 0 | 0 |
| 13 | 0 | 1 | 0 |
| 16 | 0 | 0 | 0 |
| 18 | 0 | 1.5 | 0 |
| 19 | 0 | 0.666667 | 0 |
| 21 | 1.66667 | 2 | 0.333333 |
| 22 | 0 | 1 | 0 |
| 23 | 0.25 | 1.25 | 0 |
| 24 | 0 | 0 | 0 |
| 25 | 0 | 0.666667 | 0 |
| 26 | 1 | 1.33333 | 0 |
| 27 | 0 | 1 | 0 |
| 28 | 0 | 0.333333 | 0 |
| 29 | 0.1 | 1.2 | 0 |
| 30 | 0 | 0.421053 | 0 |
| 31 | 0.0588235 | 0.647059 | 0 |
| 32 | 0.0454545 | 0.818182 | 0 |
| 33 | 0.0833333 | 0.75 | 0.0416667 |
| 34 | 0 | 0.615385 | 0.0769231 |
| 35 | 0 | 0.411765 | 0 |
| 36 | 0.0434783 | 0.782609 | 0 |
| 37 | 0 | 0.47619 | 0.047619 |
| 38 | 0.0357143 | 1.07143 | 0 |
| 39 | 0 | 0.733333 | 0 |
| 40 | 0.0689655 | 0.551724 | 0.103448 |
| 41 | 0.05 | 0.55 | 0.05 |
| 42 | 0 | 0.555556 | 0 |
| 43 | 0 | 0.466667 | 0.133333 |
| 44 | 0 | 0.947368 | 0 |
| 45 | 0 | 0.642857 | 0 |
| 46 | 0.0555556 | 0.777778 | 0.111111 |
| 47 | 0.0555556 | 0.722222 | 0.222222 |
| 48 | 0.0833333 | 0.5 | 0.333333 |
| 49 | 0.0909091 | 0.545455 | 0.454545 |
| 50 | 0.2 | 1 | 0.2 |
| 51 | 0.5 | 0.75 | 0 |
| 52 | 0 | 0.166667 | 0.166667 |
| 53 | 0 | 0 | 0 |
| 54 | 0 | 0.181818 | 0 |
| 55 | 0 | 0.12 | 0 |
| 56 | 0 | 0.153846 | 0 |
| 57 | 0 | 0.5 | 0 |
| 58 | 0 | 0.5 | 0.25 |
| 59 | 0 | 0.4 | 0 |
| 60 | 0 | 0.25 | 0 |
| 61 | 0.142857 | 0.571429 | 0.142857 |
| 62 | 0.142857 | 1 | 0 |
| 63 | 0.4 | 0.4 | 0.266667 |
| 64 | 0.0769231 | 1.11538 | 0.115385 |
| 65 | 0.0769231 | 0.423077 | 0.0769231 |
| 66 | 0.0681818 | 0.568182 | 0 |
| 67 | 0.0588235 | 0.607843 | 0.0196078 |
| 68 | 0.0447761 | 0.641791 | 0 |
| 69 | 0.0153846 | 0.476923 | 0.0615385 |
| 70 | 0.0588235 | 0.504202 | 0.0840336 |
| 71 | 0.0588235 | 0.308824 | 0.0661765 |
| 72 | 0.0405405 | 0.358108 | 0.0878378 |
| 73 | 0.0372671 | 0.546584 | 0.0372671 |
| 74 | 0.03125 | 0.325 | 0.0625 |
| 75 | 0.0588235 | 0.470588 | 0.0490196 |
| 76 | 0.0503145 | 0.383648 | 0.0125786 |
| 77 | 0.0733591 | 0.397683 | 0.015444 |
| 78 | 0.0851064 | 0.439716 | 0.0212766 |
| 79 | 0.0553746 | 0.358306 | 0.0325733 |
| 80 | 0.0424837 | 0.428105 | 0.0163399 |
| 81 | 0.0176322 | 0.518892 | 0.0125945 |
| 82 | 0.0388693 | 0.402827 | 0.0176678 |
| 83 | 0.0396825 | 0.376984 | 0.0238095 |
| 84 | 0.0446809 | 0.397872 | 0.00638298 |
| 85 | 0.0706714 | 0.436396 | 0.0388693 |
| 86 | 0.0347682 | 0.32947 | 0.0165563 |
| 87 | 0.0342577 | 0.446982 | 0.0179445 |
| 88 | 0.0752864 | 0.407529 | 0.01473 |
| 89 | 0.0610022 | 0.361656 | 0.0130719 |
| 90 | 0.04814 | 0.295405 | 0.02407 |
| 91 | 0.0456554 | 0.324006 | 0.0250368 |
| 92 | 0.0255183 | 0.317384 | 0.0414673 |
| 93 | 0.041411 | 0.343558 | 0.00920245 |
| 94 | 0.0378788 | 0.30303 | 0.00757576 |
| 95 | 0.0434783 | 0.293478 | 0.0124224 |
| 96 | 0.0332717 | 0.308688 | 0.00554529 |
| 97 | 0.0662768 | 0.267057 | 0.0175439 |
| 98 | 0.0252101 | 0.283313 | 0.00960384 |
| 99 | 0.0497738 | 0.312217 | 0.00603318 |
| 100 | 0.031339 | 0.316239 | 0.0185185 |
| 101 | 0.0402098 | 0.283217 | 0.0122378 |
| 102 | 0.018 | 0.256 | 0.014 |
| 103 | 0.0209302 | 0.255814 | 0.0209302 |
| 104 | 0.0233051 | 0.245763 | 0.00847458 |
| 105 | 0.0243902 | 0.251524 | 0.0259146 |
| 106 | 0.0336283 | 0.281416 | 0.00707965 |
| 107 | 0.0261569 | 0.289738 | 0.0301811 |
| 108 | 0.0371901 | 0.289256 | 0.0371901 |
| 109 | 0.0469314 | 0.238267 | 0.0252708 |
| 110 | 0.0418118 | 0.310105 | 0.0209059 |
| 111 | 0.0200803 | 0.273092 | 0.0281124 |
| 112 | 0.0268456 | 0.238255 | 0.0201342 |
| 113 | 0.0166205 | 0.204986 | 0.0221607 |
| 114 | 0.0175439 | 0.266667 | 0.0175439 |
| 115 | 0.0310345 | 0.255172 | 0.0344828 |
| 116 | 0.00699301 | 0.384615 | 0.041958 |
| 117 | 0.0384615 | 0.307692 | 0.00769231 |
| 118 | 0.018018 | 0.333333 | 0.027027 |
| 119 | 0.0160428 | 0.262032 | 0.0160428 |
| 120 | 0.041958 | 0.244755 | 0 |
| 121 | 0.0347222 | 0.333333 | 0.0138889 |
| 122 | 0.0169492 | 0.338983 | 0 |
| 123 | 0.010989 | 0.230769 | 0.010989 |
| 124 | 0.0144928 | 0.434783 | 0.0434783 |
| 125 | 0.0210526 | 0.252632 | 0.0736842 |
| 126 | 0.025641 | 0.264957 | 0.0512821 |
| 127 | 0.0921053 | 0.381579 | 0.0526316 |
| 128 | 0.032967 | 0.384615 | 0.021978 |
| 129 | 0.0365854 | 0.280488 | 0.0121951 |
| 130 | 0.0181818 | 0.345455 | 0.0181818 |
| 131 | 0 | 0.151515 | 0.030303 |
| 132 | 0.0652174 | 0.26087 | 0 |
| 133 | 0 | 0.36 | 0.02 |
| 134 | 0 | 0.0625 | 0 |
| 135 | 0 | 0.0588235 | 0 |
| 136 | 0 | 0.2 | 0 |
| 137 | 0 | 0.153846 | 0 |
| 138 | 0 | 0.333333 | 0.166667 |
| 139 | 0 | 0.2 | 0 |
| 140 | 0 | 0 | 0 |
| 142 | 0 | 1 | 0 |

**Figure S4.** The average number of changes (y-axes) observed among samples of different ages (x-axes). Changes are shown on four plots according to their starting base, and colored within each plot according to their resulting base.

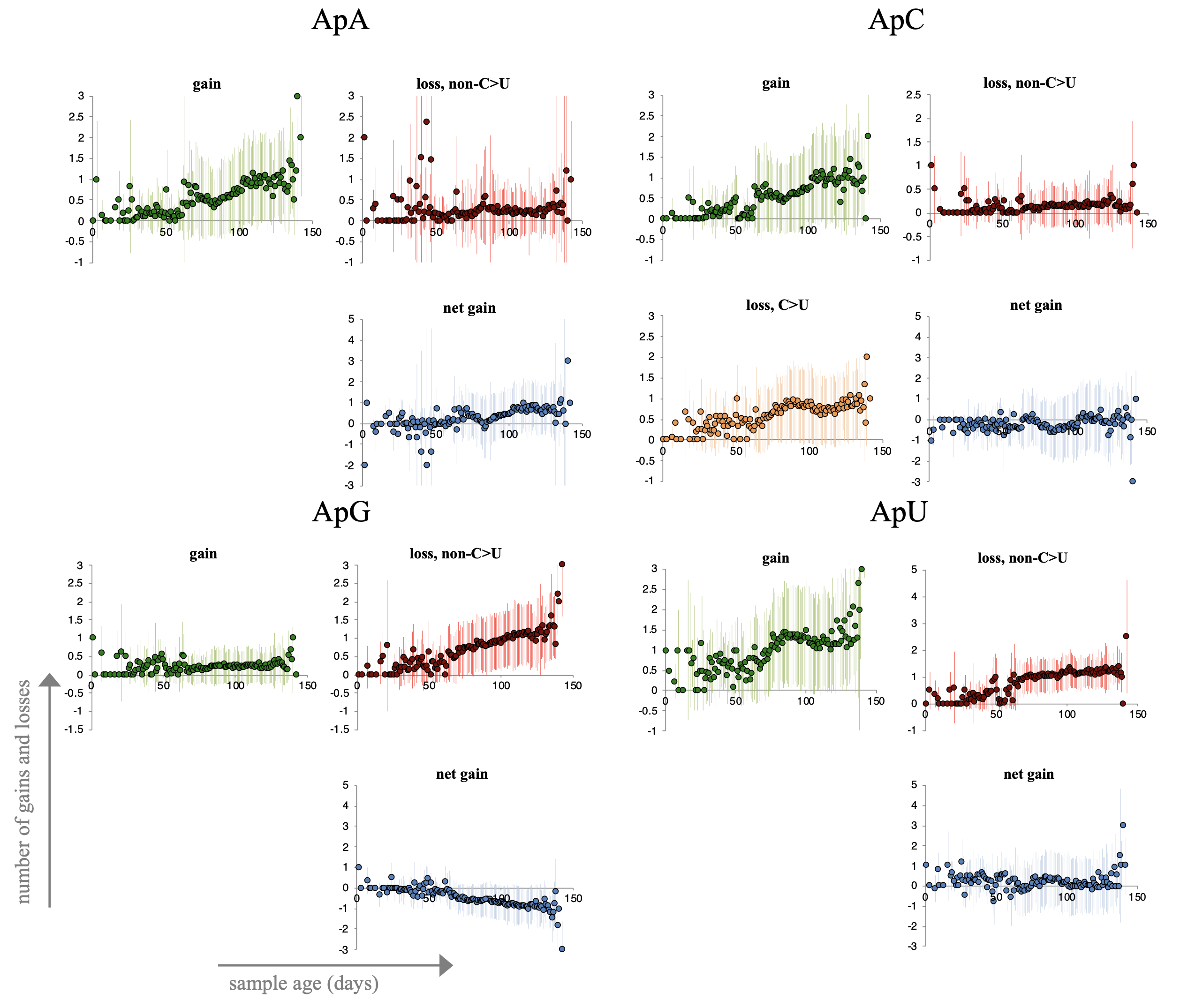

**Figure S5.** Number of AN dinucleotide gains and losses over time. For all dinucleotides the number of gains and losses over time are shown. Numbers are calculated as averages across all samples from a given day (x-axes) and shown with plus/minus one standard deviation. Green plots show gains, red plots show losses through other changes than C-to-U, orange plots (when applicable) show losses through C-to-U changes. Finally, blue plots show net gains simply calculated as gains minus losses

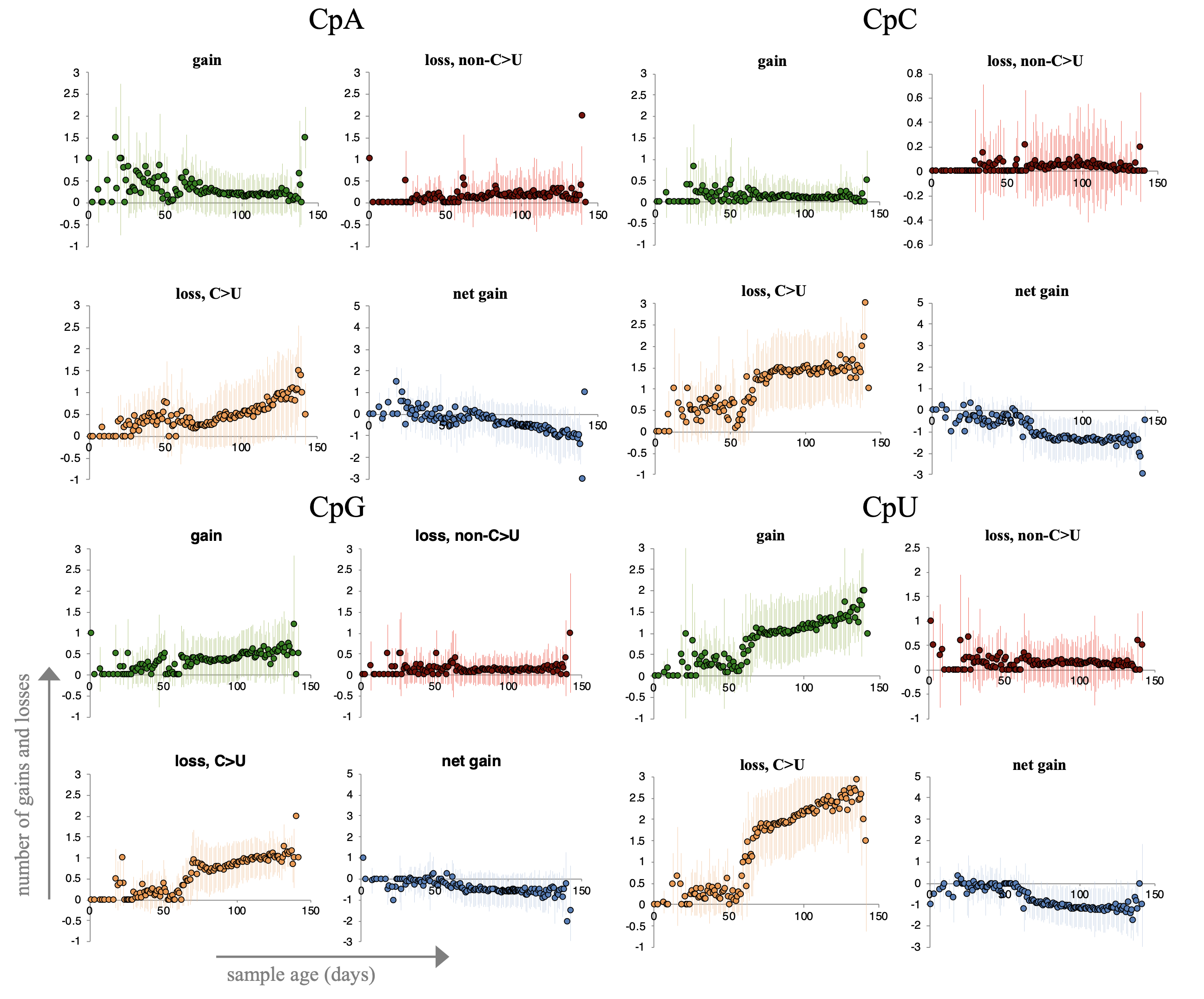

**Figure S5 (cont.).** Number of CN dinucleotide gains and losses over time.

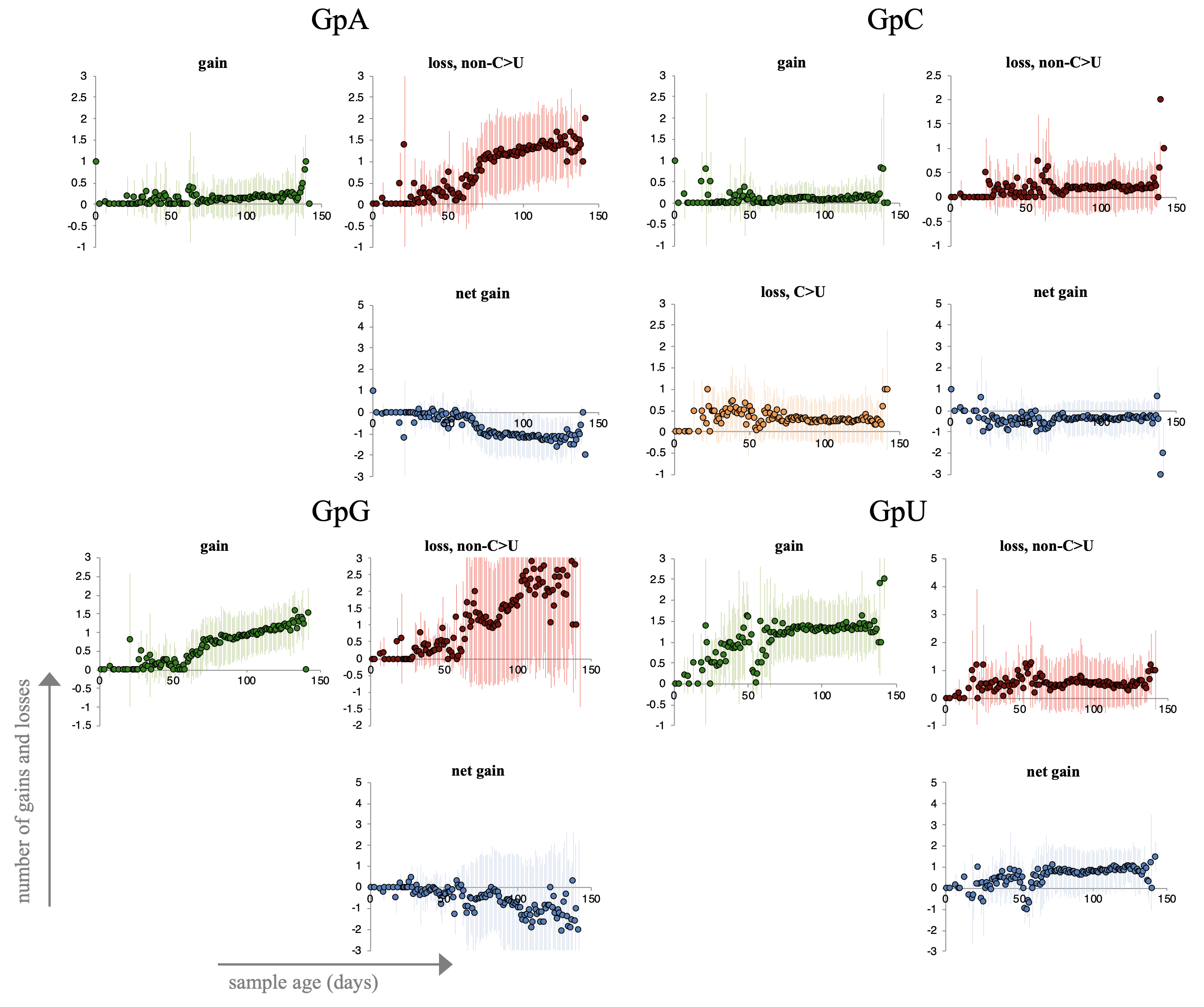

**Figure S5 (cont.).** Number of GN dinucleotide gains and losses over time.

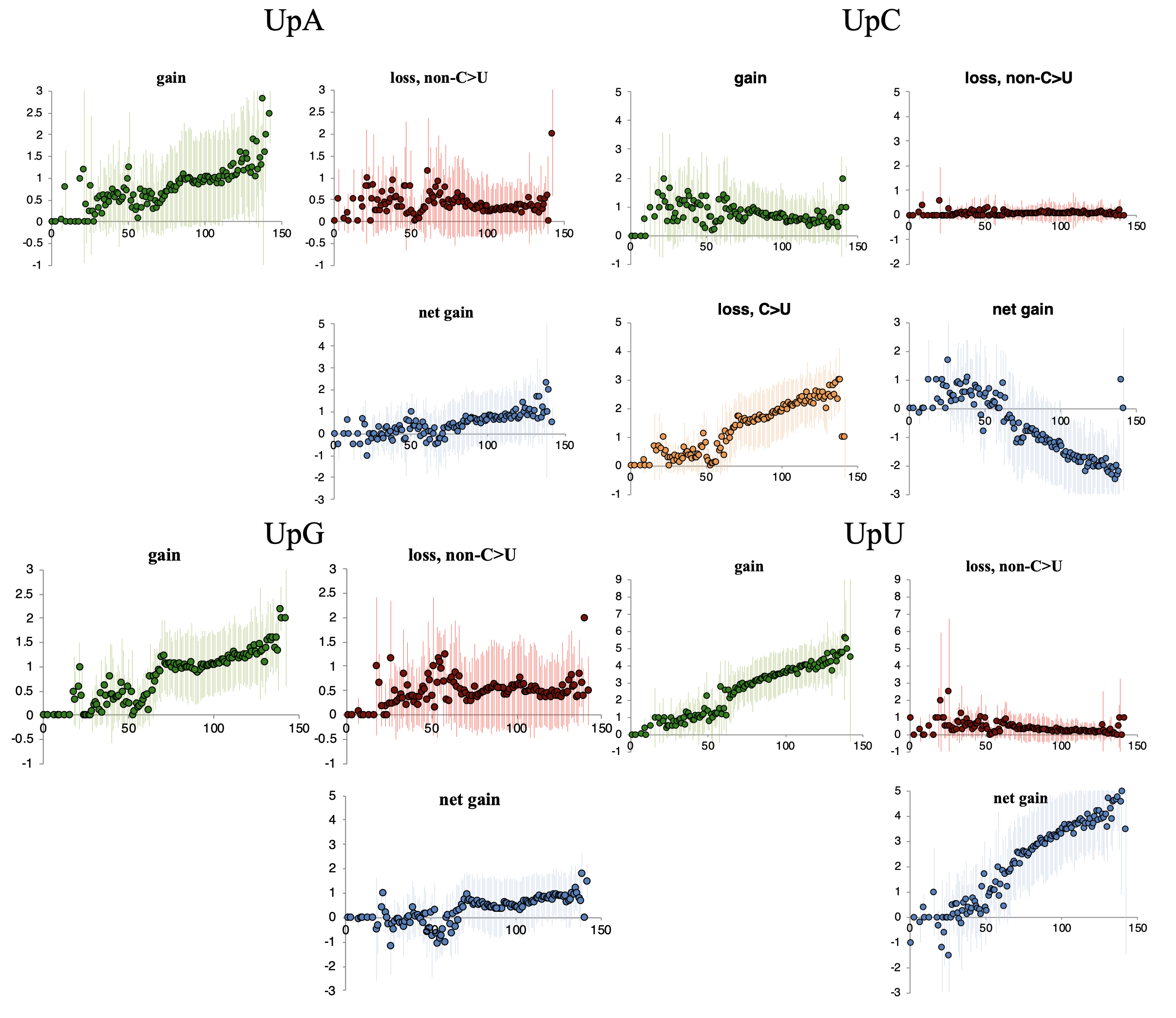
 **Figure S5 (cont.).** Number of UN dinucleotide gains and losses over time.

position in genome

fraction of triplets

number of triplets

Fraction of triplets that have

UCG-to-UUG changes

(right y-axis)

UCG triplets in reference genome

(left y-axis)

|  | triplets with UCG>UUG | UCG triplets |
| --- | --- | --- |
| 1000 | 0.4 | 15 |
| 2000 | 0.6 | 5 |
| 3000 | 0.2 | 5 |
| 4000 | 0.666667 | 3 |
| 5000 | 1 | 1 |
| 6000 |  | 0 |
| 7000 | 0.666667 | 3 |
| 8000 | 1 | 2 |
| 9000 | 0 | 3 |
| 10000 | 1 | 2 |
| 11000 | 0.333333 | 3 |
| 12000 | 0 | 1 |
| 13000 |  | 0 |
| 14000 | 0.2 | 5 |
| 15000 | 0.333333 | 3 |
| 16000 | 0.25 | 4 |
| 17000 | 0.5 | 2 |
| 18000 | 0.25 | 4 |
| 19000 | 0.333333 | 3 |
| 20000 | 0.5 | 2 |
| 21000 | 0.25 | 4 |
| 22000 | 0 | 2 |
| 23000 | 0.5 | 2 |
| 24000 | 0 | 1 |
| 25000 | 0 | 1 |
| 26000 | 0.25 | 4 |
| 27000 | 0.181818 | 11 |
| 28000 | 0.25 | 4 |
| 29000 | 0.3 | 10 |
| 30000 | 0.625 | 8 |

**Figure S6.**

Number of UCG triplets in 1kb bins in the reference genome (line with blue squares).

The fraction of UCG triplets for which a change to UUG is observed in any sample is shown as light blue bars.

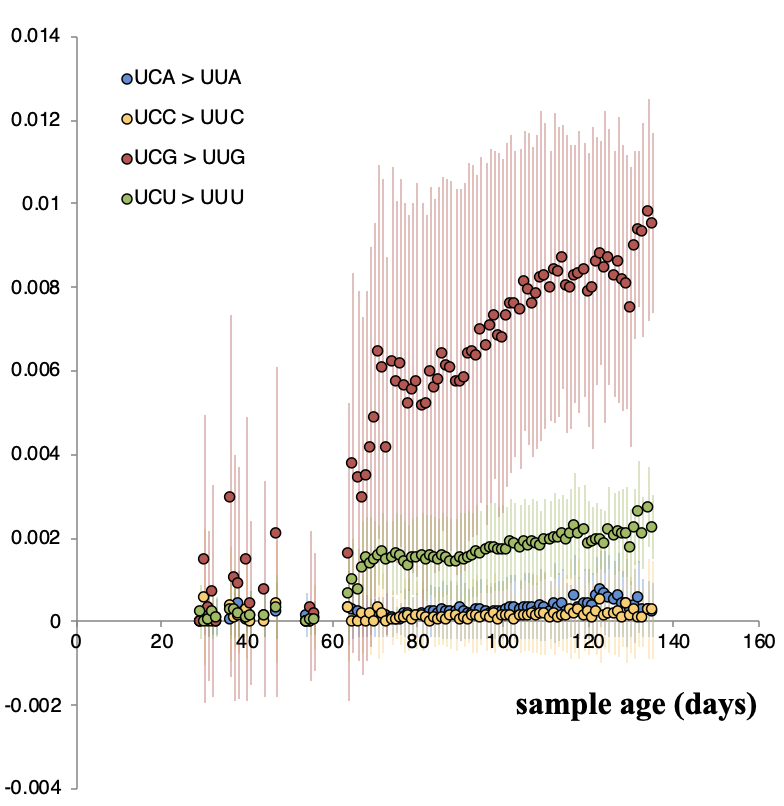

**Figure S7.** The fraction of UC[A/C/G/U] to UU[A/C/G/U] changes over time. The fraction of trinucleotide changes relative to the reference genome are shown as circles for each sampling day (x-axis). Error bars denote plus/minus one standard deviation.

```
(+) 5’- CGN - 3’
        |||
(-) 3’- GCN - 5’
```

```
(+) 5’- CAN – 3’
        |||
(-) 3’- GUN – 5’
```

Cytosine deamination

```
(+) 5’- NGA - 3’
        |||
(-) 3’- NCU - 5’
```

```
(+) 5’- NAA – 3’
        |||
(-) 3’- NUU – 5’
```

Cytosine deamination

|  | CGA>CAA | CGC>CAC | CGG>CAG | CGT>CAT |
| --- | --- | --- | --- | --- |
| 32 | 0 | 0 | 0 | 0 |
| 33 | 0 | 0.000859107 | 0 | 0 |
| 36 | 0 | 0.000448229 | 0 | 0.000254259 |
| 37 | 0 | 0 | 0 | 0.000139237 |
| 38 | 0.00112782 | 0.000736377 | 0 | 0 |
| 40 | 0 | 0 | 0 | 0.000201654 |
| 41 | 0 | 0.000515464 | 0 | 0.000584795 |
| 54 | 0 | 0 | 0 | 0 |
| 55 | 0 | 0 | 0 | 0 |
| 56 | 0 | 0 | 0 | 0 |
| 64 | 0.000809717 | 0 | 0.000506073 | 0 |
| 65 | 0 | 0 | 0 | 0 |
| 66 | 0 | 0 | 0 | 0 |
| 67 | 0 | 0 | 0.000257998 | 0 |
| 68 | 0 | 0 | 0.000196386 | 0 |
| 69 | 0 | 0 | 0 | 0 |
| 70 | 0 | 0 | 0.000221141 | 0 |
| 71 | 7.73994e-05 | 0 | 0.000193498 | 0 |
| 72 | 7.11238e-05 | 0 | 0 | 0 |
| 73 | 0 | 6.40328e-05 | 0 | 0 |
| 74 | 0 | 0.000128866 | 0 | 0 |
| 75 | 0 | 0 | 0 | 0 |
| 76 | 6.62032e-05 | 0.000324191 | 8.27541e-05 | 0 |
| 77 | 0 | 0 | 0.000152408 | 2.2579e-05 |
| 78 | 0 | 3.65577e-05 | 4.66592e-05 | 0 |
| 79 | 0 | 0.000100742 | 4.28596e-05 | 0 |
| 80 | 0 | 0.000101071 | 0 | 0 |
| 81 | 7.95439e-05 | 0.000155808 | 0 | 1.47304e-05 |
| 82 | 0 | 0.000182143 | 0.000139483 | 4.13283e-05 |
| 83 | 0 | 8.18197e-05 | 5.22139e-05 | 0 |
| 84 | 2.23964e-05 | 6.58039e-05 | 0 | 4.97698e-05 |
| 85 | 0 | 0.0001275 | 4.64943e-05 | 1.03321e-05 |
| 86 | 1.74277e-05 | 6.82734e-05 | 0 | 8.71384e-05 |
| 87 | 3.43436e-05 | 0.000117724 | 2.14648e-05 | 0 |
| 88 | 1.7228e-05 | 0.00011811 | 2.1535e-05 | 2.87134e-05 |
| 89 | 2.29331e-05 | 6.73809e-05 | 2.86664e-05 | 2.54813e-05 |
| 90 | 2.30335e-05 | 0.000112793 | 5.75838e-05 | 2.55928e-05 |
| 91 | 1.55027e-05 | 0.000212562 | 0.000155027 | 3.44504e-05 |
| 92 | 3.35768e-05 | 8.22112e-05 | 2.09855e-05 | 3.73075e-05 |
| 93 | 3.22893e-05 | 7.90589e-05 | 0 | 2.69078e-05 |
| 94 | 1.99362e-05 | 7.81006e-05 | 4.98405e-05 | 5.53783e-05 |
| 95 | 8.17261e-05 | 9.60492e-05 | 8.17261e-05 | 9.98874e-05 |
| 96 | 0 | 9.52798e-05 | 0 | 5.40476e-05 |
| 97 | 2.05191e-05 | 0.000140672 | 0 | 2.2799e-05 |
| 98 | 3.79099e-05 | 9.90087e-05 | 1.57958e-05 | 2.10611e-05 |
| 99 | 3.17536e-05 | 0.000124396 | 9.923e-05 | 2.64613e-05 |
| 100 | 1.49948e-05 | 0.000205598 | 1.87434e-05 | 4.16521e-05 |
| 101 | 1.84026e-05 | 0.000162209 | 2.30033e-05 | 0 |
| 102 | 0 | 2.06186e-05 | 2.63158e-05 | 0.000116959 |
| 103 | 2.44798e-05 | 0 | 0 | 5.43996e-05 |
| 104 | 6.69045e-05 | 2.18417e-05 | 0 | 2.47795e-05 |
| 105 | 0 | 0.000110008 | 4.01155e-05 | 3.56583e-05 |
| 106 | 3.72613e-05 | 0.000109479 | 2.32883e-05 | 3.10511e-05 |
| 107 | 2.11797e-05 | 6.2229e-05 | 7.94239e-05 | 2.3533e-05 |
| 108 | 4.34972e-05 | 8.52006e-05 | 5.43715e-05 | 9.66604e-05 |
| 109 | 0 | 0.00014887 | 0 | 2.11117e-05 |
| 110 | 0 | 3.59208e-05 | 4.58463e-05 | 8.15046e-05 |
| 111 | 0 | 8.28054e-05 | 0 | 4.69715e-05 |
| 112 | 0 | 6.91898e-05 | 4.4154e-05 | 0 |
| 113 | 2.91588e-05 | 0.000171345 | 7.28969e-05 | 4.85979e-05 |
| 114 | 0 | 0.000108519 | 0.000138504 | 4.10383e-05 |
| 115 | 0 | 0.000213295 | 4.53721e-05 | 8.06614e-05 |
| 116 | 0 | 0 | 0 | 0.000163579 |
| 117 | 0 | 0 | 0 | 0 |
| 118 | 0 | 0 | 0.000237079 | 5.26843e-05 |
| 119 | 0 | 0.00011026 | 0 | 9.38174e-05 |
| 120 | 7.36106e-05 | 0 | 9.20132e-05 | 4.08948e-05 |
| 121 | 0 | 0.000143184 | 9.13743e-05 | 8.12216e-05 |
| 122 | 0 | 8.73668e-05 | 0 | 4.95589e-05 |
| 123 | 0 | 0 | 0 | 0 |
| 124 | 0.000305111 | 0.00014941 | 0 | 8.47529e-05 |
| 125 | 0.000221607 | 0.000108519 | 0 | 6.15574e-05 |
| 126 | 8.99685e-05 | 8.81135e-05 | 0.000112461 | 0 |
| 127 | 0.000277008 | 0.000135648 | 0.00034626 | 0 |
| 128 | 0.000115674 | 0 | 0.000144592 | 0 |
| 129 | 0.00012837 | 0 | 0.000160462 | 0 |
| 130 | 0 | 0.000187441 | 0.000239234 | 0.000212653 |
| 131 | 0 | 0 | 0 | 0 |
| 132 | 0 | 0 | 0.000286041 | 0 |
| 133 | 0 | 0.000206186 | 0 | 0 |
| 134 | 0 | 0 | 0 | 0.000182749 |
| 135 | 0 | 0 | 0 | 0.000171999 |

|  | CGA>CAA | AGA>AAA | GGA>GAA | TGA>TAA |
| --- | --- | --- | --- | --- |
| 32 | 0 | 0 | 0 | 0 |
| 33 | 0 | 0 | 0 | 0 |
| 36 | 0 | 0 | 0 | 0 |
| 37 | 0 | 3.93546e-05 | 0 | 0 |
| 38 | 0.00112782 | 0 | 0.000126646 | 0 |
| 40 | 0 | 0 | 0 | 0 |
| 41 | 0 | 0 | 0 | 0 |
| 54 | 0 | 7.51315e-05 | 0 | 0.0001443 |
| 55 | 0 | 0 | 0 | 6.34921e-05 |
| 56 | 0 | 0 | 9.09256e-05 | 8.14001e-05 |
| 64 | 0.000809717 | 0 | 0 | 0 |
| 65 | 0 | 0 | 0 | 0 |
| 66 | 0 | 0 | 8.05932e-05 | 3.6075e-05 |
| 67 | 0 | 0 | 6.95314e-05 | 0 |
| 68 | 0 | 0 | 0 | 0 |
| 69 | 0 | 0 | 0 | 2.442e-05 |
| 70 | 0 | 1.38899e-05 | 0 | 0 |
| 71 | 7.73994e-05 | 0 | 0 | 0 |
| 72 | 7.11238e-05 | 0 | 7.18804e-05 | 0 |
| 73 | 0 | 1.02664e-05 | 0 | 0 |
| 74 | 0 | 3.09917e-05 | 0 | 0 |
| 75 | 0 | 0 | 0 | 0 |
| 76 | 6.62032e-05 | 0 | 0 | 0 |
| 77 | 0 | 0 | 1.36915e-05 | 0 |
| 78 | 0 | 0 | 0 | 5.62873e-06 |
| 79 | 0 | 5.38401e-06 | 2.31016e-05 | 5.17036e-06 |
| 80 | 0 | 5.40161e-06 | 1.15886e-05 | 0 |
| 81 | 7.95439e-05 | 4.16346e-06 | 1.78645e-05 | 1.99912e-05 |
| 82 | 0 | 1.16812e-05 | 0 | 1.12177e-05 |
| 83 | 0 | 1.31182e-05 | 0 | 6.29882e-06 |
| 84 | 2.23964e-05 | 7.03359e-06 | 4.52694e-05 | 3.37724e-06 |
| 85 | 0 | 1.16812e-05 | 2.50608e-05 | 1.40221e-05 |
| 86 | 1.74277e-05 | 1.9156e-05 | 2.34841e-05 | 7.88395e-06 |
| 87 | 3.43436e-05 | 2.6964e-06 | 1.15697e-05 | 1.03576e-05 |
| 88 | 1.7228e-05 | 1.35261e-05 | 3.48226e-05 | 1.81851e-05 |
| 89 | 2.29331e-05 | 7.20215e-06 | 3.09028e-05 | 6.91635e-06 |
| 90 | 2.30335e-05 | 2.53178e-05 | 3.10381e-05 | 1.04199e-05 |
| 91 | 1.55027e-05 | 1.94744e-05 | 3.13352e-05 | 2.3377e-05 |
| 92 | 3.35768e-05 | 2.10895e-05 | 2.82783e-05 | 2.27842e-05 |
| 93 | 3.22893e-05 | 1.01404e-05 | 1.08776e-05 | 1.46071e-05 |
| 94 | 1.99362e-05 | 1.56524e-05 | 1.34322e-05 | 9.01876e-06 |
| 95 | 8.17261e-05 | 1.79662e-05 | 2.20255e-05 | 1.23238e-05 |
| 96 | 0 | 1.52763e-05 | 1.31094e-05 | 8.80204e-06 |
| 97 | 2.05191e-05 | 9.66604e-06 | 2.76499e-05 | 1.23766e-05 |
| 98 | 3.79099e-05 | 1.98426e-05 | 2.55421e-05 | 2.47718e-05 |
| 99 | 3.17536e-05 | 2.74236e-05 | 3.744e-05 | 9.57648e-06 |
| 100 | 1.49948e-05 | 1.41273e-05 | 1.51543e-05 | 1.13056e-05 |
| 101 | 1.84026e-05 | 2.02277e-05 | 3.71968e-05 | 5.55001e-06 |
| 102 | 0 | 3.30579e-05 | 2.12766e-05 | 1.5873e-05 |
| 103 | 2.44798e-05 | 2.69076e-05 | 2.47402e-05 | 7.3828e-06 |
| 104 | 6.69045e-05 | 2.45132e-05 | 3.00517e-05 | 1.68146e-05 |
| 105 | 0 | 2.77162e-05 | 2.70282e-05 | 2.41967e-06 |
| 106 | 3.72613e-05 | 2.34038e-05 | 1.25526e-05 | 8.42815e-06 |
| 107 | 2.11797e-05 | 2.32802e-05 | 2.1405e-05 | 9.5813e-06 |
| 108 | 4.34972e-05 | 6.83013e-06 | 2.93066e-05 | 6.5591e-06 |
| 109 | 0 | 4.17698e-05 | 2.56036e-05 | 5.73033e-06 |
| 110 | 0 | 4.03145e-05 | 3.70672e-05 | 5.53067e-06 |
| 111 | 0 | 3.31906e-05 | 2.84827e-05 | 6.37471e-06 |
| 112 | 0 | 2.21865e-05 | 3.5699e-05 | 5.32652e-06 |
| 113 | 2.91588e-05 | 4.12078e-05 | 2.9469e-05 | 0 |
| 114 | 0 | 5.21966e-05 | 8.70972e-05 | 3.34169e-05 |
| 115 | 0 | 2.84981e-05 | 4.89117e-05 | 0 |
| 116 | 0 | 1.15587e-05 | 7.43937e-05 | 1.11e-05 |
| 117 | 0 | 6.35728e-05 | 8.18331e-05 | 0 |
| 118 | 0 | 5.95637e-05 | 3.19468e-05 | 1.43e-05 |
| 119 | 0 | 4.4195e-05 | 1.89631e-05 | 0 |
| 120 | 7.36106e-05 | 3.46761e-05 | 0.000173585 | 1.11e-05 |
| 121 | 0 | 1.14784e-05 | 0.000197006 | 6.61376e-05 |
| 122 | 0 | 0 | 3.00517e-05 | 1.34517e-05 |
| 123 | 0 | 3.63273e-05 | 3.89681e-05 | 1.74429e-05 |
| 124 | 0.000305111 | 2.3955e-05 | 0.000102785 | 2.30044e-05 |
| 125 | 0.000221607 | 6.95955e-05 | 0 | 0 |
| 126 | 8.99685e-05 | 4.23819e-05 | 3.03085e-05 | 0 |
| 127 | 0.000277008 | 4.34972e-05 | 9.33184e-05 | 0 |
| 128 | 0.000115674 | 9.08183e-05 | 0.000311745 | 0 |
| 129 | 0.00012837 | 4.03145e-05 | 4.32451e-05 | 3.87147e-05 |
| 130 | 0 | 0 | 0 | 2.886e-05 |
| 131 | 0 | 0 | 0 | 0 |
| 132 | 0 | 7.18649e-05 | 0 | 0 |
| 133 | 0 | 0 | 7.0922e-05 | 0 |
| 134 | 0 | 0 | 0 | 4.96032e-05 |
| 135 | 0 | 4.86145e-05 | 0 | 0 |

**Figure S8.** Cytosine deamination of NCG and UCN triplets on the viral negative strand.

If APOBEC deaminates UCG to UUG on the negative strand this will manifest itself as a CGA-to-CAA change on the positive strand.

On the top plot, the progression of CG(N)-to-CA(N) changes are shown, testing the base upstream of deaminated CpG. The progression of (N)GA-to-(N)AA is shown on the bottom plot, testing the base downstream of deaminated UpC.

For all plotted changes, no apparent progression over time is observed, suggesting that the action of APOBEC on the negative strand of SARS-CoV-2 is limited compared to the positive strand.

**Figure S9.** The inverse relationship of C>U frequency and the number of samples included. Down-sizing the number of samples leads to higher C>U frequency (0.1 on the x-axis means 10% of the total samples), suggesting that as more and more samples are included the prominent C>U changes happen as independent events at identical positions (hence only counted once in this analysis). As this saturation is expected to be lower for other, less abundant changes this will result in an apparent drop in the fraction of changes constituted by C>U. The blue dots are the individual simulated data sets, and the boxes denote their distribution.

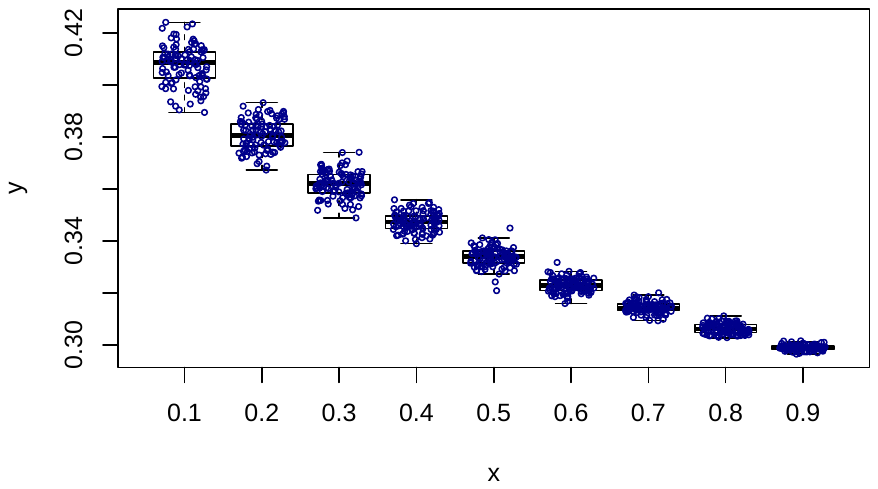

sub-fraction of samples

C>U frequency

**Table S2.** The most enriched motifs undergoing C>U changes.

| **C>U** | | | |
| --- | --- | --- | --- |
| motif | motifs with C-to-U change/total number of motifs | odds-ratio | uncorrected p-value |
| UCG | 72/113 | 1.30 | 0.0012387 |
| UUCG | 27/39 | 1.41 | 0.0088244 |
| CG | 257/439 | 1.19 | 4.56E-05 |
| CGG | 55/76 | 1.47 | 3.19E-05 |
| CGGA | 13/16 | 1.65 | 0.0088056 |
| CGGU | 22/29 | 1.54 | 0.0030566 |
| GUGC | 75/113 | 1.35 | 0.0001577 |
| UGCU | 148/255 | 1.18 | 0.0026032 |
| CUA | 307/561 | 1.11 | 0.0044054 |
| UUAC | 137/241 | 1.16 | 0.0096264 |
| UGAC | 87/137 | 1.29 | 0.0004834 |
| CAAC | 112/194 | 1.18 | 0.0098096 |
| UAAC | 88/135 | 1.33 | 0.00012 |

**Supplementary References**

Atkinson NJ, Witteveldt J, Evans DJ, Simmonds P. 2014. The influence of CpG and UpA dinucleotide frequencies on RNA virus replication and characterization of the innate cellular pathways underlying virus attenuation and enhanced replication. Nucleic Acids Res. *42*(7):4527–4545. doi:10.1093/nar/gku075.

Digard P, Lee HM, Sharp C, Grey F, Gaunt E. 2020. Intra-genome variability in the dinucleotide composition of SARS-CoV-2. Virus Evol. *6*(2). doi:10.1093/ve/veaa057.

Elbe S, Buckland-Merrett G. 2017. Data, disease and diplomacy: GISAID’s innovative contribution to global health. Glob Challenges. *1*(1):33–46. doi:10.1002/gch2.1018.

Di Gioacchino A, Sulc P, Komarova A V., Greenbaum BD, Monasson R, Cocco S. 2020. The Heterogeneous Landscape and Early Evolution of Pathogen-Associated CpG Dinucleotides in SARS-CoV-2. SSRN Electron J. doi:10.2139/ssrn.3611280.

Greenbaum BD, Levine AJ, Bhanot G, Rabadan R. 2008. Patterns of evolution and host gene mimicry in influenza and other RNA viruses. PLoS Pathog. *4*(6). doi:10.1371/journal.ppat.1000079.

Katoh K, Standley DM. 2013. MAFFT multiple sequence alignment software version 7: Improvements in performance and usability. Mol Biol Evol. *30*(4):772–780. doi:10.1093/molbev/mst010.

Lloyd DCF, Sykes P. 1978. An introduction to business games. Ind Commer Train. *10*(1):11–18. doi:10.1108/eb003648.

Lorenz R, Bernhart SH, Höner zu Siederdissen C, Tafer H, Flamm C, Stadler PF, Hofacker IL. 2011. ViennaRNA Package 2.0. Algorithms Mol Biol. *6*(1). doi:10.1186/1748-7188-6-26.

Poulain F, Lejeune N, Willemart K, Gillet NA. 2020. Footprint of the host restriction factors APOBEC3 on the genome of human viruses. PLoS Pathog. *16*(8). doi:10.1371/JOURNAL.PPAT.1008718.

Rice AM, Castillo Morales A, Ho AT, Mordstein C, Mühlhausen S, Watson S, Cano L, Young B, Kudla G, Hurst LD. 2020. Evidence for Strong Mutation Bias toward, and Selection against, U Content in SARS-CoV-2: Implications for Vaccine Design. Mol Biol Evol. doi:10.1093/molbev/msaa188.

Simmonds P, Xia W, Baillie JK, McKinnon K. 2013. Modelling mutational and selection pressures on dinucleotides in eukaryotic phyla -selection against CpG and UpA in cytoplasmically expressed RNA and in RNA viruses. BMC Genomics. *14*(1). doi:10.1186/1471-2164-14-610.

Stern A, Yeh M Te, Zinger T, Smith M, Wright C, Ling G, Nielsen R, Macadam A, Andino R. 2017. The Evolutionary Pathway to Virulence of an RNA Virus. Cell. *169*(1):35-46.e19. doi:10.1016/j.cell.2017.03.013.

Takata MA, Gonçalves-Carneiro D, Zang TM, Soll SJ, York A, Blanco-Melo D, Bieniasz PD. 2017. CG dinucleotide suppression enables antiviral defence targeting non-self RNA. Nature. *550*(7674):124–127. doi:10.1038/nature24039.

Wang Y, Mao JM, Wang GD, Luo ZP, Yang L, Yao Q, Chen KP. 2020. Human SARS-CoV-2 has evolved to reduce CG dinucleotide in its open reading frames. Sci Rep. *10*(1):12331. doi:10.1038/s41598-020-69342-y.

Wei Y, Silke J, Aris P, Xia X. 2020. Coronavirus genomes carry the signatures of their habitats. bioRxiv. doi:10.1101/2020.06.13.149591.

Woo PCY, Wong BHL, Huang Y, Lau SKP, Yuen KY. 2007. Cytosine deamination and selection of CpG suppressed clones are the two major independent biological forces that shape codon usage bias in coronaviruses. Virology. *369*(2):431–442. doi:10.1016/j.virol.2007.08.010.

Xia X. 2020. Extreme genomic CpG deficiency in SARS-CoV-2 and evasion of host antiviral defense. Mol Biol Evol. *37*(9):2699–2705. doi:10.1093/molbev/msaa094.

Yu X, Yu Y, Liu B, Luo K, Kong W, Mao P, Yu XF. 2003. Induction of APOBEC3G Ubiquitination and Degradation by an HIV-1 Vif-Cul5-SCF Complex. Science (80- ). *302*(5647):1056–1060. doi:10.1126/science.1089591.

Zhang J, Kang J, Liu M, Han B, Li L, He Y, Yi Z, Chen L. 2020 Jan 1. Multi-site co-mutations and 5’UTR CpG immunity escape drive the evolution of SARS-CoV-2. bioRxiv.:2020.07.21.213405. doi:10.1101/2020.07.21.213405.
